# ALPHA: A High Throughput System for Quantifying Growth In Aquatic Plants

**DOI:** 10.1101/2024.07.30.605820

**Authors:** Kassidy A. Robinson, Victoria Augoustides, Tanaka Madenyika, Ryan C. Sartor

## Abstract

The need for more sustainable agricultural systems is becoming increasingly apparent. The global demand for agricultural products — food, feed, fuel and fiber — will continue to increase as the global population continues to grow. This challenge is compounded by climate change. Not only does a changing climate make it difficult to maintain stable yields but current agricultural systems are a major source of greenhouse gas emissions and continue to drive the problem further. Therefore, future agricultural systems must not only increase production but also significantly decrease negative environmental impacts. One approach to addressing this is to begin breeding and cultivating new plant species that have fundamental sustainability advantages over our existing crops. The Lemnaceae, a.k.a duckweeds, are one such species that have potential to increase output and reduce the negative environmental impacts of agricultural production. Herein we describe the Automated Lab-scale PHenotyping Apparatus, ALPHA, for high-throughput phenotyping of Lemnaceae. ALPHA is being used for selective breeding of one species, *Lemna gibba*, toward the goal of creating a new crop for use in sustainable agricultural systems. ALPHA can be used on many small aquatic plant species to assess growth rates in different environmental conditions. A proof of principle use case is demonstrated where ALPHA is used to determine saltwater tolerance of 6 different varieties of *L. gibba*.

## Introduction

Lemnaceae, commonly known as “duckweed”, is a family of floating aquatic plants characterized by their rapid growth rate, simple anatomical structure and the ability to tolerate and accumulate high nutrient levels [Walsh, 2021; Ziegler, 2023]. As the global population continues to grow in the coming decades, significant changes to our agricultural systems will be necessary. We will require sustainable and scalable production of food, feed, fuel and fiber. Wild plant species have the potential to contribute to this transition significantly; by leveraging natural traits and adaptations, we can breed/develop novel crops that are better suited to meet the developing needs of human society. Lemnaceae species can be utilized for both production of agricultural products and remediation of agricultural production due to three key traits: 1) They grow well on municipal and animal waste and can efficiently uptake nutrients from wastewater [Porath, 1982; Cheng, 2002; Mohedano, 2012; Cui, 2015; Zhou, 2021]. 2) They grow fast and are high yielding. With yield potential as high as 20 tons/acre/year, they can produce several times more biomass per acre than existing crops [Ziegler, 2015; Cui, 2015]. 3) They produce high quality, nutritious biomass, with protein as high as 35% of dry mass and starch as high as 30% of dry mass [Appenroth, 2017]. These traits make Lemnaceae an ideal candidate for reducing our agricultural sector’s carbon footprint on multiple fronts. Through the phytoremediation of agricultural waste, Lemnaceae can tackle nutrient pollution while simultaneously producing high-quality biomass that can be used for animal feed or bioproducts [Walsh, 2021].

The Lemna Improvement and Functional Transcriptomics (LIFT) Lab at North Carolina State University focuses on selectively breeding *Lemna gibba*, a species of Lemnaceae, as a platform for sustainable agricultural systems. We refer to *L. gibba* simply as “Lemna”. Building on top of the aforementioned natural traits, the LIFT lab has identified two major traits to target for improvement: Density tolerance and temperature tolerance. Improvement of these traits will dramatically increase both yields and yield stability, making *L. gibba* a viable agricultural crop and a tool for sustainable agriculture.

Successful selective breeding relies on accurate and reproducible phenotyping at multiple stages. We are working to implement a series of successive phenotyping systems, which progress from lab-scale high-throughput to open-pond field trials. This Lemna “breeding funnel” allows for a progressive tradeoff where high throughput is prioritized at the beginning and more accurate, larger-scale trait evaluation is prioritized in the final system.

Lemna’s small size affords lab-scale phenotyping in tightly regulated, high throughput conditions, and herein, we describe a phenotyping platform that accomplishes the first level of high-throughput phenotyping. To efficiently measure growth rates of *L. gibba*, the LIFT Lab developed the Automated Lab-scale Phenotyping Apparatus (ALPHA). This high-throughput imaging system allows for rapid, consistent imaging of Lemna cultures and allows for growth in many desired conditions. ALPHA utilizes an automated image analysis program, created with PlantCV [Gehan, 2017], to accurately detect plants within an image and produce valuable phenotypic data. This data, specifically plant area, facilitates the measurement of growth rates on an array of *L. gibba* lines. When breeding, this phenotypic data is then used for selection of the highest performing lines in the desired environmental conditions.

Prior work towards image-based phenotyping of duckweed has either been designed for high resolution microscopy-based imaging at the cost of low-throughput [Cox, 2022] or designed with limited growth space not suitable for larger species and requiring a large, custom growth chamber [Kose, 2023]. ALPHA’s novel design uses 20 × 150 mm culture tubes as opposed to tissue culture plates, to provide ample surface area and depth for aquatic plant growth. Culture tubes are also less prone to spilling, contamination, are reusable, more sustainable, and generally easier to handle compared to well plates. In addition, the system implements a side-view during imaging that shows the depth of the duckweed mat, as well as a barcode. ALPHA’s design uncouples the plant growth from the imaging, allowing the system to be scaled to meet demand without requiring large growth chambers or grow rooms. The remainder of this paper describes the hardware and software of ALPHA, our image analysis pipeline, and a proof-of principle example of how ALPHA was used to determine salinity tolerance in 6 *L. gibba* varieties.

## Materials and Methods

### Source Code, 3D models and Data

All source code used in the phenotyping system, 3D models for printed parts and data generated for this study are available in the ALPHA Github repository.

### L. gibba Growth

All *L. gibba* varieties screened in ALPHA are cultured in axenic conditions. *L. gibba* cultures are imaged in 20 × 150 mm culture tubes with clear KimKap caps to allow for ventilation while maintaining a sterile environment. Total culture volume is 10 ml of sterile liquid media, and any liquid media formulation of appropriate optical density can be used for imaging. The tubes are housed in custom 3D-printed growth racks (**Fig. 1B**) that easily load into ALPHA and stand upright in growth chambers when not imaging (**Fig. 1D**). The tubes are randomized within the racks during experiments, and the racks are rotated around the incubator daily to minimize positional effects within the growth chambers.

**Figure 1.**
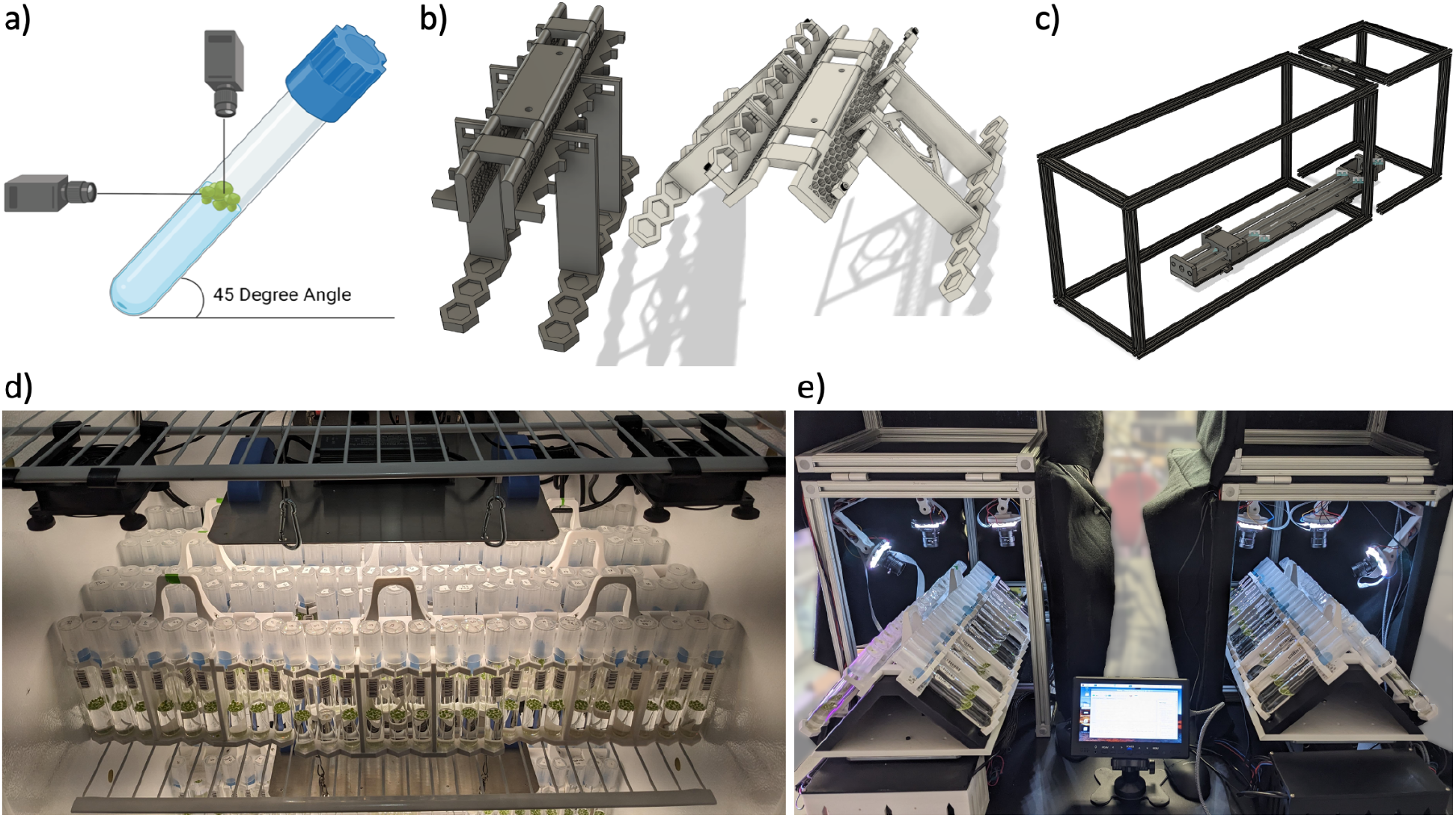
Components of the ALPHA system. a) Dual cameras for top and side image capture. b) 3D printed tube rack designed for imaging at a 45° angle. c) Layout of imaging system frame and actuator. d) Tube racks in the growth chamber in the upright position. e) Tube racks on dual imagers in the 45° position.

### Custom 3D-printed Culture Tube Racks

Culture tubes are maintained and transported in custom 3D-printed racks consisting of a main body, two wings and a handle (**Fig. 1B**). The rack assembly accommodates both 90 degree growth conditions in a “closed” conformation, and 45 degree imaging conditions in an “open” conformation due to three hinges at the center of the main rack body, one longer central hinge and two shorter flanking hinges. Each wing can support 9 culture tubes with a midbody contact point to stabilize the tube during transport, and a load-bearing platform. To account for sub-millimeter manufacturing variability in tube diameter, tube holsters are hexagonally-shaped, and adjustable via a small gap to allow flexure in the plastic. We chose to print our racks in polylactic acid (PLA) due to its flexibility, accessibility, and cost-effectiveness though other materials are likely also sufficient.

### ALPHA Imaging System

The outer structure of the imaging system is constructed from T-slotted extruded aluminum that forms a cage around the inner components (**Fig. 1C**), with one hinged section at the front to allow for easy opening and closure. A black felt cover blocks ambient light while imaging. Each rack sits on a metal platform, the “actuator carriage”, and is ferried in the imaging system on a 69 cm motorized ball screw linear actuator. The actuator is rigidly secured to the bottom of the outer structure, and driven with a stepper motor. A custom 3D-printed platform is fixed atop the actuator carriage which acts as the mount for the test tube racks. The triangular shape of the platform holds each wing of the racks at a 45 degree angle, allowing the top-view cameras to capture the surface area of the tube and the side-view cameras to capture the barcode and depth of the mat (**Fig. 1E**). Cut outs in the platform secure the racks in place while imaging and eliminate any shifting. The track length of the actuator carriage is regulated by two optical limit switch sensors that detect and define the track endpoints to prevent the actuator carriage from moving too far in either direction. The entire actuator system is controlled by an Arduino microcontroller. This setup allows for precise and reproducible placement of the tubes for imaging.

A Raspberry Pi 4 Model B is used as the master controller for the entire imaging system. It directly controls the cameras, and interfaces with the Arduino microcontroller. The Raspberry Pi is equipped with an Arducam multi camera board allowing four high quality (12 Megapixel) cameras to be controlled from a single Raspberry Pi. Each camera is mounted on an articulated 3D printed camera arm [Hamkers, 2018] and attached to the aluminum structure. The design of the camera arms allows the angle and position of the cameras to be easily adjusted and fine-tuned. Individual LED array light sources for each camera are mounted on the end of each arm. ALPHA uses two top-view cameras with 8 mm lenses to capture the surface area of the tube, as well as two side-view cameras with 6 mm lenses to capture the depth of the duckweed mat and a barcode containing the unique identifier for the tube. The cameras are controlled using PiCamera2 [Plowman, 2023], an open source Python library for Raspberry Pi cameras.

In addition to controlling the position of the tube rack with the stepper motor, the Arduino also controls the LED arrays that act as lights sources for each camera. Source code can be found in “Alpha_arduino_code.ino” in the Github repository. The camera lights are RGB NeoPixel Rings, each with 24 individually programmable LEDs where the RGB values and intensity of each LED can be individually adjusted to optimize the light quality for imaging. Typically we use only half of the LEDs on each NeoPixel Ring to reduce glare from the tubes and have lowered the blue value after noticing the sample images had a blue tint. The Adafruit “NeoPixel” Arduino library [Adafruit, 2022] is used for controlling these lights.

Serial communication – the process of sending information bit by bit over a communication channel – is a fundamental part of ALPHA. More specifically, bidirectional serial communication is used to facilitate communication between the Arduino and Raspberry Pi. A custom software interface was developed using the Arduino C-based language and the Python PySerial [Liechti, 2020] package on the Raspberry Pi. This interface allows the Raspberry Pi to act as a master controller, sending commands to the Arduino via Python. This increases the functionality and efficiency of the system, allowing for precise coordination of the lights, cameras, stepper motor, and two optical limit switch sensors.

### Image Acquisition Cycle

ALPHA is operated by executing a Python script on the Raspberry Pi. Source code can be found in “plant_phenotyping.py” in the Github repository. At the beginning of the script, serial communication between the Raspberry Pi and the Arduino is established. To begin the automated imaging, two racks of tubes (36 samples) are loaded onto the imaging platform. The Raspberry Pi instructs the Arduino to return the platform to the starting position, moving it backward until it reaches the first optical limit switch. Now that the platform position is reset, the imaging can begin. The platform is then moved to the first position for imaging which is calibrated to the center of a group of three culture tubes. Next, for each camera, the lights are turned on one at a time and an image is captured. Each image contains 3 samples/tubes and is tagged with the current timestamp and the camera identification number (1-4). Once all four photos have been taken for the first position, the platform moves to the next position, and continues the same process until all tubes have been imaged. At this point, the platform’s position is reset and returns to the starting point. During imaging, the images are stored locally on the Raspberry Pi. At the end of each day, all photos taken within the last 24 hours are automatically uploaded to the lab’s Google Drive using a Linux cron job.

### PlantCV Image Analysis

A Python workflow using the open source PlantCV software [Gehan, 2017] was developed to quantify phenotypic data from the images and output the data to a CSV file. Source code can be found in “plant_processing.py” in the Github repository. A series of functions are used to detect the plant surface area within the top-view images and the depth of the duckweed mat as well as a barcode, from the side-view images. For the top-view images, the image is first converted from RGB color space to LAB color space and split by the green-magenta channel using plantcv.rgb2gray_lab() (**Fig. 2b**). The resulting grayscale image is then used to create a binary mask, in which the detected plant in the image is white, and the rest is black (**Fig. 2c**). The pcv.threshold.binary() function is used to generate this mask by determining which pixels are included in the white mask and which are included in the black background. Optimizing this threshold is crucial for accurately detecting plant area. Next, plantcv.fill() is used to filter out noise within the mask (**Fig. 2d**). Adjusting this threshold is especially important when imaging smaller duckweed species, as they can go undetected if the threshold is set too high. The mask is then overlaid with the image using plantcv.find_objects(), and the duckweed colonies are detected as objects (**Fig. 2e)**. This composite image is then split into three separate sections, or regions of interest (ROI), using the plantcv.roi.multi() function (see **Fig. 2f**, blue circles). Each ROI represents one of 3 samples in the image. Finally, objects within each ROI are grouped together using plantcv.object_composition(), then the total plant surface area for each tube is calculated with plantcv.analyze_object() (**Fig. 2g-i**). The resulting data is outputted to a JSON file.

**Figure 2.**
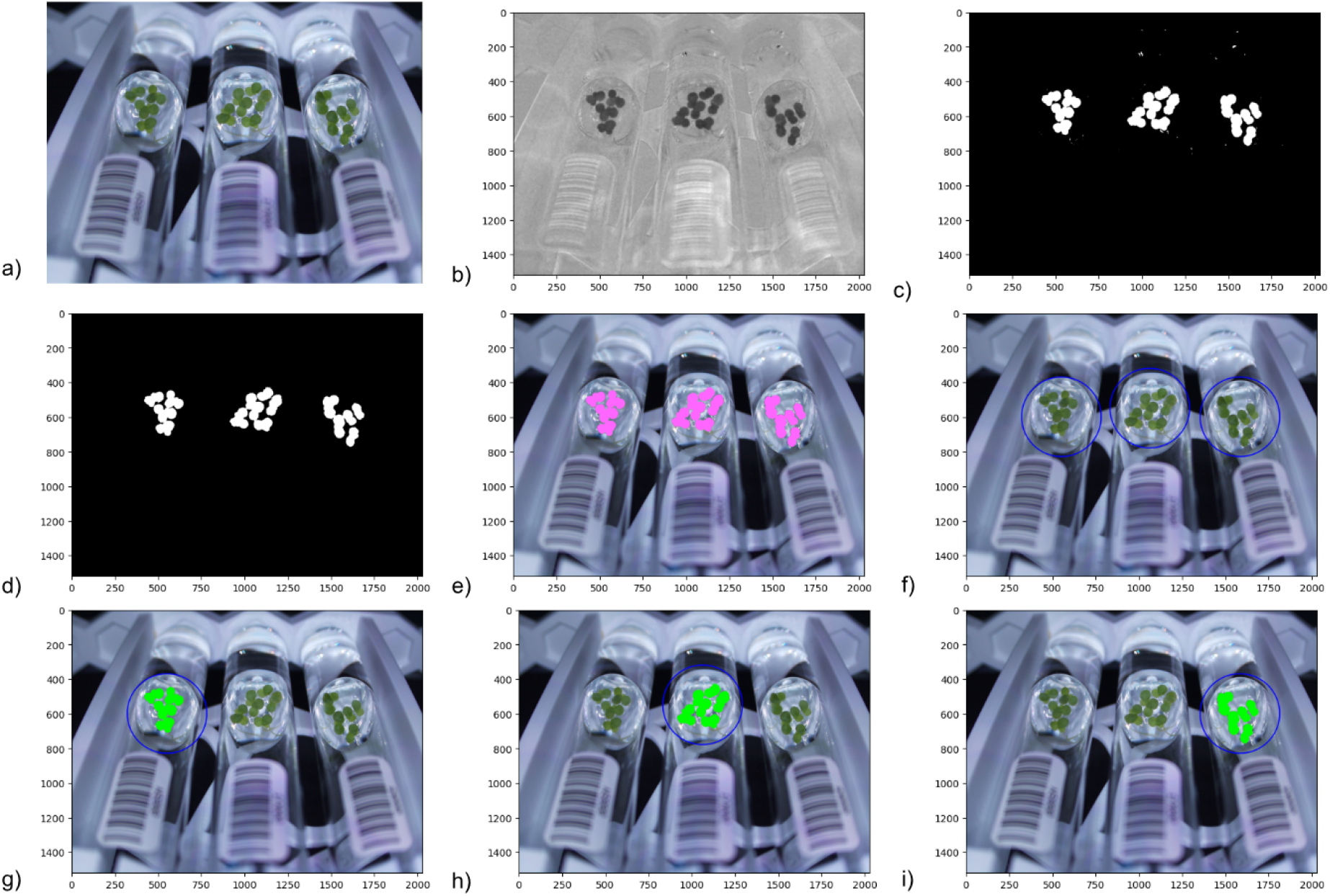
a) Top-view image. b) LAB color space image split by the green-magenta channel. c) Binary mask of the green detected in the image. d) Binary mask with noise filtered out. e) Objects detected (in pink) by overlaying the binary mask with the original image. f) Circular regions of interest (ROIs) for each tube. g-i) Objects detected (in green) within each ROI.

For the side-view images, a similar workflow was created (**Fig 3a-d**). However, instead of using the PlantCV plantcv.analyze_object() to determine the depth of the floating plant mat, we wrote a function to more accurately measure the depth. Our function uses the binary mask of the plant area, reads through each row of pixels, and calculates the number of white pixels in each row. If the total number of white pixels for a row reaches a set threshold, that row is included in the height measurement. This threshold ensures that the depth is only calculated if the culture is dense enough to grow across the entire tube, and reduces inaccurate spikes in height by eliminating rows that are below the threshold.

**Figure 3.**
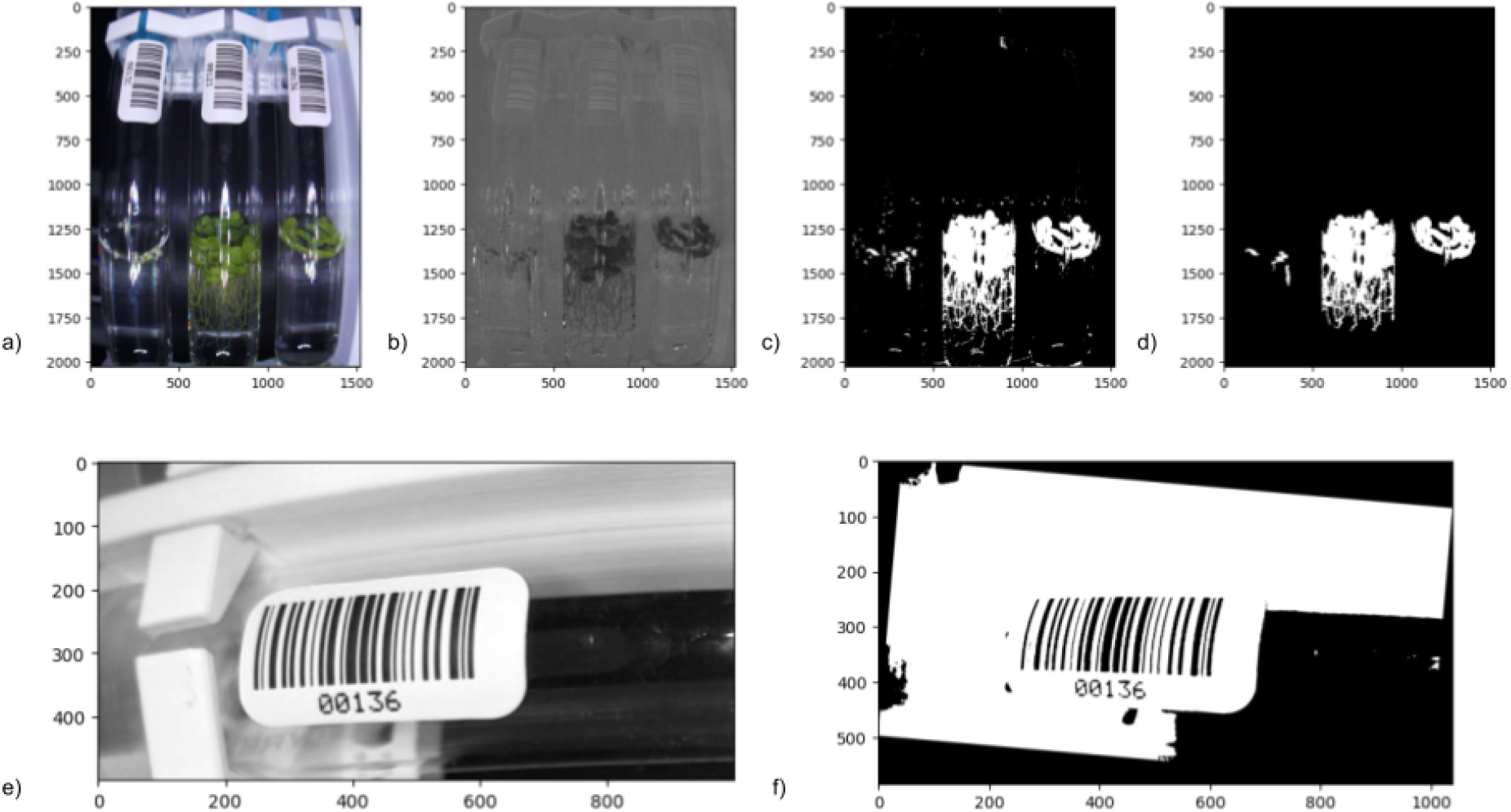
a) Rotated side-view image. b) LAB color space image split by the green-magenta channel. c) Binary mask of the green detected in the image. d) Binary mask with noise filtered out. e) Side-view image, converted to grayscale and cropped to select for the barcode. f) Final binary mask that Pyzbar was able to detect the barcode from.

Pyzbar [Hudson, 2022], a Python package that utilizes the open source ZBar library, was used to detect and read barcodes from the side-view images. The barcodes contain unique sample identifiers and are used to keep track of all samples during a phenotyping run. For each image, the image is cropped to focus on the tube’s barcode and Pyzbar is used to read the barcode. Since each image can have slight variations in exposure and rotations of the labels, the software often has trouble detecting the barcode. An algorithm is used to cycle the image through thresholds and degrees of rotation until pyzbar.decode() is able to detect the barcode. This significantly increases the likelihood of barcodes being accurately read.

The data from these side-view images is added to the same JSON file as the data from the top-view images. After the program has finished running, the JSON file is converted to a more user-friendly CSV format.

### Generating Growth Curves

The following analysis was done using the R statistical programming language and open source software packages (R core, 2020). The source code can be found in “ALPHA_GrowthCurves.R” in the Github repository. This code requires data output from the quantification pipeline “PlantCV_Output_Salinity.csv” and the barcode map “Barcode_Sample_Map.csv”. Both are also available in the Github repository. Before the phenotypic data can be analyzed, the CSV files are first manipulated using a series of functions in R to match the data from the top-view images with the data from the corresponding side-view images. All data from each sample is combined into a singular row containing the image name, camera number, timestamp, area, barcode, and depth. Each phenotyping run will span several weeks and can have dozens of images for each sample. Using the unique IDs, read from the barcodes, the data from each tube are matched and sorted by timestamp to create growth curves for each sample. These growth curves are the primary product of the phenotyping system and are used to assess the response of each individual sample to the experimental conditions.

### Salinity Tolerance Trial

Six different *L. gibba* varieties were acquired from the Rutgers Duckweed Stock Cooperative (RDSC) [“Rutgers Duckweed Stock Cooperative, 2024]. The strains are hereafter referred to by the country or region of origin. The RDSC accessions 7805, 9352, 7263, 7641, 8678 and 8682 are referred to as “France”, “Germany”, “Greece”, “Israel”, “Kashmir” and “Saudi” respectively. Each of these varieties were grown under 6 different salinity concentrations. Schenk and Hildebrandt basil salts were used at ½ concentration (1.6 g/Liter) to provide nutrients to all samples. Instant Ocean® synthetic sea salt was added to increase salinity. This mix contains 26.28% sodium by total mass [Dickman, 2002]. Six concentrations were formulated to give the following final concentrations of added sodium: 0mM, 50mM, 100mM, 200mM, 300mM, and 400mM. All six varieties were tested in all six concentrations with six replicates each for a total of 216 cultures. A single, small colony was added to each tube to start the culture. Plants were grown at a constant 25°C with 14 hours light and 10 hours dark under an average light intensity of 800 μmol/m^2^/s. Tubes were randomly assigned to racks and the rack positions inside the growth chamber were rotated daily. Each rack was imaged 5 days a week, for 4 weeks, resulting in 20 days of data.

### Salinity Tolerance Analysis

All analysis was done using the R statistical programming language and open source software packages (R core, 2020). Source code can be found in “SaltAnalysis.R” in the Github repository. The growth curves must first be generated with “ALPHA_GrowthCurves.R”. Briefly, growth curves were derived from each sample as described above. Samples were filtered out that did not contain plant area values of 1000 pixels for at least 15 consecutive days to remove samples that did not grow or died.

Relative growth rates (RGR) were calculated along each curve using a sliding 10-day window. For each RGR, a model fitness is also returned. The maximum RGR with at least 0.8 adjusted R-squared fitness is used as the RGR for each sample. At 200mM concentrations and above, no plants grew so zero values for RGR were substituted in for the 200mM samples and all higher concentrations were removed. Next, the drc package (Ritz, 2015) was used to fit dose-response curves to the data using a 5-factor model. The EC50 is defined as the salt concentration at which plants grow at 50% of the optimal growth rate (0mM added salt). The EC_50_ for each plant variety was determined using the fitted model equations and solving for 50% of the 0mM RGR value. The “boot” R package (Canty & Ripley, 2024) was used to estimate the standard deviation of EC_50_s. All data was plotted using the “ggplot2” (Wickham, 2016) and “scales” (Wickham, 2020) R packages.

## Results

Replacing fresh water with sea water in agriculture can lead to significant water savings. Here, as a proof of concept, ALPHA was used to assess the sea water tolerance of six varieties of *L. gibba*. Instant Ocean® synthetic sea salt was used to increase the salinity of normal culture media. Six different salinity concentrations were tested (0mM, 50mM, 100mM, 200mM, 300mM, and 400mM). These refer to the amount of additional sodium that was added using synthetic sea salt and assuming the salt mix consists of 26.28% sodium by mass [Dickman, 2002]. For reference, sea water contains ∼ 480mM of sodium [Johnson, 1979]. Plant cultures were grown for 4 weeks and imaged 5 days per week.

The images were processed using our PlantCV pipeline, and analyzed using R as described in the methods. The analysis showed that the 200mM, 300mM, and 400mM concentrations were too high for the *L. gibba* cultures to survive. The RGR values for the 200mM samples were set to zero and the higher concentrations were removed from the analysis. Response curves for all 6 varieties are shown in **figure 4**. Growth rates vary between varieties at low salt concentrations but are similar at the highest survivable concentration of 100mM.

**Figure 4.**
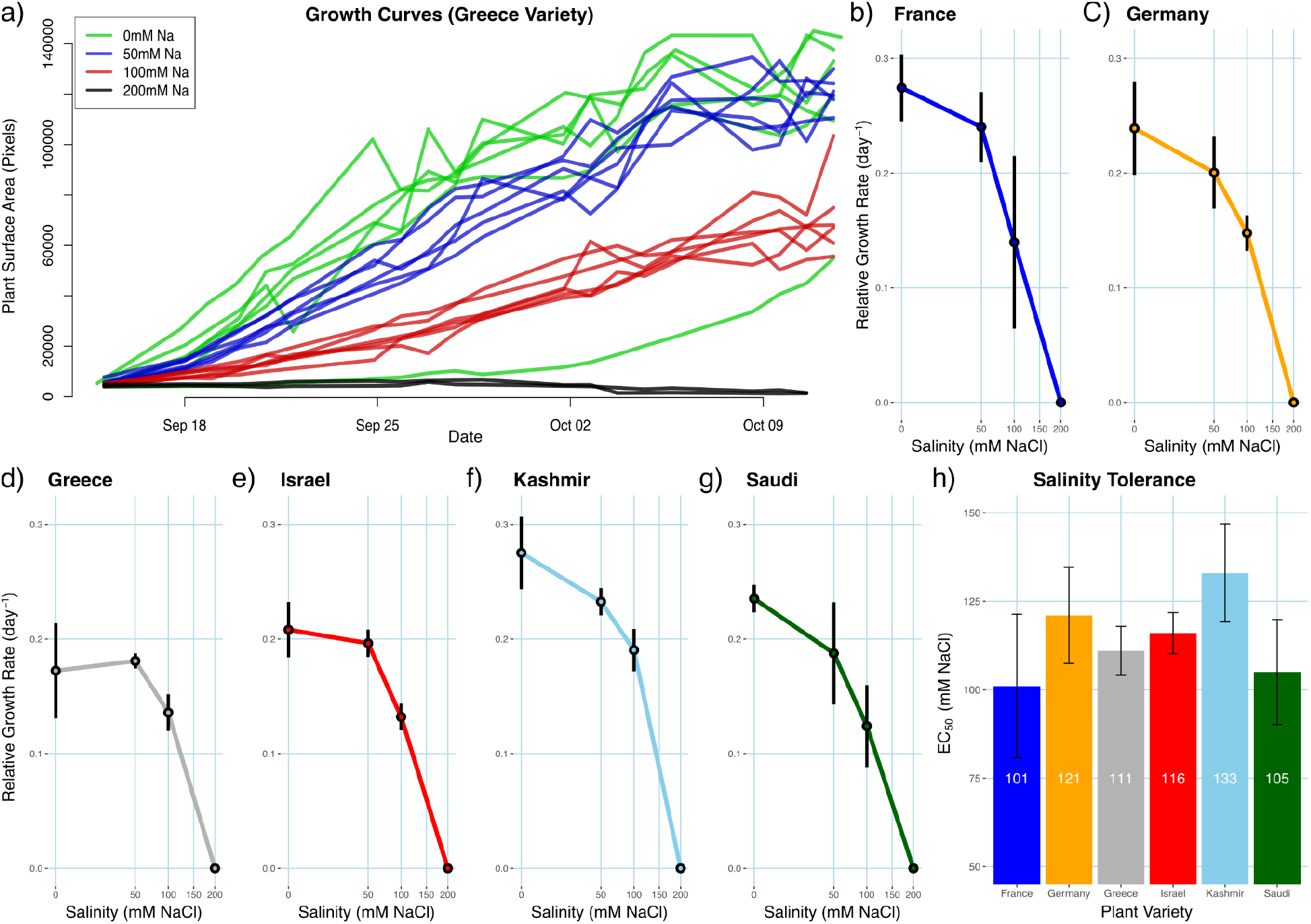
*L. gibba* salinity tolerance. a) Representative growth curves for the Greece variety in 4 different salinity concentrations, n = 6 for each condition. b-g) Line plots of relative growth rates vs. Salinity for 6 different *L*.*gibba* varieties. h) Barplot showing the EC_50_ (salinity tolerance) for all 6 *L. gibba* varieties.

Salinity tolerance was determined by calculating the EC_50_ (Effective Concentration at 50% maximum growth rate) for each variety (**Fig 4**). The most salinity tolerant variety was found to be the “Kashmir” variety (RDSC# 8678) with the lowest being the “France” variety (RDSC# 7805). While significant differences in growth rate are observed between varieties at every concentration lower than 200mM, this trial did not identify any significant differences in salt tolerances (EC_50_) between the varieties tested. The “Kashmir” variety maintained the highest relative growth rate of 0.19 day^-1^ at the 100mM concentrations of sea salt. This trial implies that this variety could potentially tolerate 20% of typical ocean salinity while maintaining 70% of its optimal growth rate.

## Conclusion

ALPHA, a system for high-throughput phenotyping of small aquatic plants, has been developed. This system is capable of determining plant growth rates in any desired media and at different temperatures or light regimes. The application of ALPHA is demonstrated by using it to determine salinity tolerance of 6 different varieties of the species *L. gibba*. The system is capable of phenotyping any small, floating aquatic species and could potentially work for submerged species as well. The system is useful for studying basic biological processes through characterization of growth rates under different environmental conditions or to measure traits for quantitative genetic studies. The system can also be used in applied breeding programs for aquatic plants as a method for phenotyping and selection using fast breeding cycles. ALPHA will be used for both purposes in the future.

While the current state of ALPHA is sufficient for use, one area of improvement is the quantification software. Currently no machine learning methods are being used with this system. Plant detection and quantification relies on traditional methods of thresholding on RGB values. A convolutional neural network based approach for object detection and segmentation may have advantages over the current method. These advanced methods are currently being prototyped with ALPHA.

The example provided shows the utility of ALPHA for assessing production potential upon partial supplementation of seawater for irrigation. Future breeding goals are going to also pursue the creation of a new crop for use in integrated animal production systems that efficiently recycles nutrients. Lemna has been shown to remediate animal waste while producing high quality biomass for the production of biofuels [Xu, 2011; Cui, 2015] and animal feed [Leng 1995; Rojas, 2014; Demman, 2022].

## Data Availability

GitHub: https://github.com/LiftLaboratory/Alpha. This repository contains all code associated with the phenotyping system and the analysis done in this manuscript. The repository also contains design details for construction of the system along with all 3D models for 3D printed parts of the system.

## Author Contributions

ALPHA was conceived and designed by RS, VA and KR.

ALPHA hardware was built by VA, KR and RS.

ALPHA software was developed by KR, RS with contributions from VA.

Salinity tolerance analysis was carried out by KR and TM.

The Manuscript was written by KR with edits from RS and VA.

